# Fast, scalable and accurate differential expression analysis for single cells

**DOI:** 10.1101/049734

**Authors:** Debarka Sengupta, Nirmala Arul Rayan, Michelle Lim, Bing Lim, Shyam Prabhakar

## Abstract

Analysis of single-cell RNA-seq data is challenging due to technical variability, high noise levels and massive sample sizes. Here, we describe a normalization technique that substantially reduces technical variability and improves the quality of downstream analyses. We also introduce a nonparametric method for detecting differentially expressed genes that scales to > 1,000 cells and is both more accurate and ~10 times faster than existing parametric approaches.

## MAIN TEXT

Single-cell RNA sequencing (scRNA-seq) technologies have revolutionized functional genomics by providing a window into heterogeneity within complex cellular ensembles.^1^ However, substantial challenges remain in analyzing the resulting data. Single-cell transcriptomes are distorted by technical biases such as RNA degradation during cell isolation and processing, variable reagent amounts, presence of cellular debris and PCR amplification bias.^2,3^ Moreover, due to the small number of molecules under investigation (< 5 transcripts per moderately expressed gene), single-cell expression estimates are inherently noisy,^2^ even in the absence of technical variability. Thus, the two most basic components of transcriptomic data analysis, normalization and differential expression (DE) analysis, are uniquely challenging for scRNAseq. In addition, the introduction of new, high-throughput technologies such as inDrop/ Drop-seq^4^ has created a pressing need for algorithms that can scale beyond a thousand cells.

The most widely used RNA-seq normalization methods, which include Fragments Per Kilobase per Million reads^5^(FPKM), size factor^6^ (DEseq algorithm) and Trimmed Mean of M-values^7^ (TMM; edgeR algorithm), were originally developed for bulk RNA-seq data. These methods assume that the number of reads for a given transcript is directly proportional to its expression level,after adjusting for technical covariates. If this assumption were true, the distribution of expression estimates (counts, FPKMs) would have a consistent shape across samples of the same type, and this is indeed frequently observed in bulk data (Fig. 1a). However, we found that the shape of the scRNA-seq expression distribution was highly variable from cell to cell (Fig. 1b,c), even in the absence of cell cycle variability (Fig. 1d). Thus, the proportionality assumption may not be appropriate for scRNA-seq normalization.

**Figure 1.**
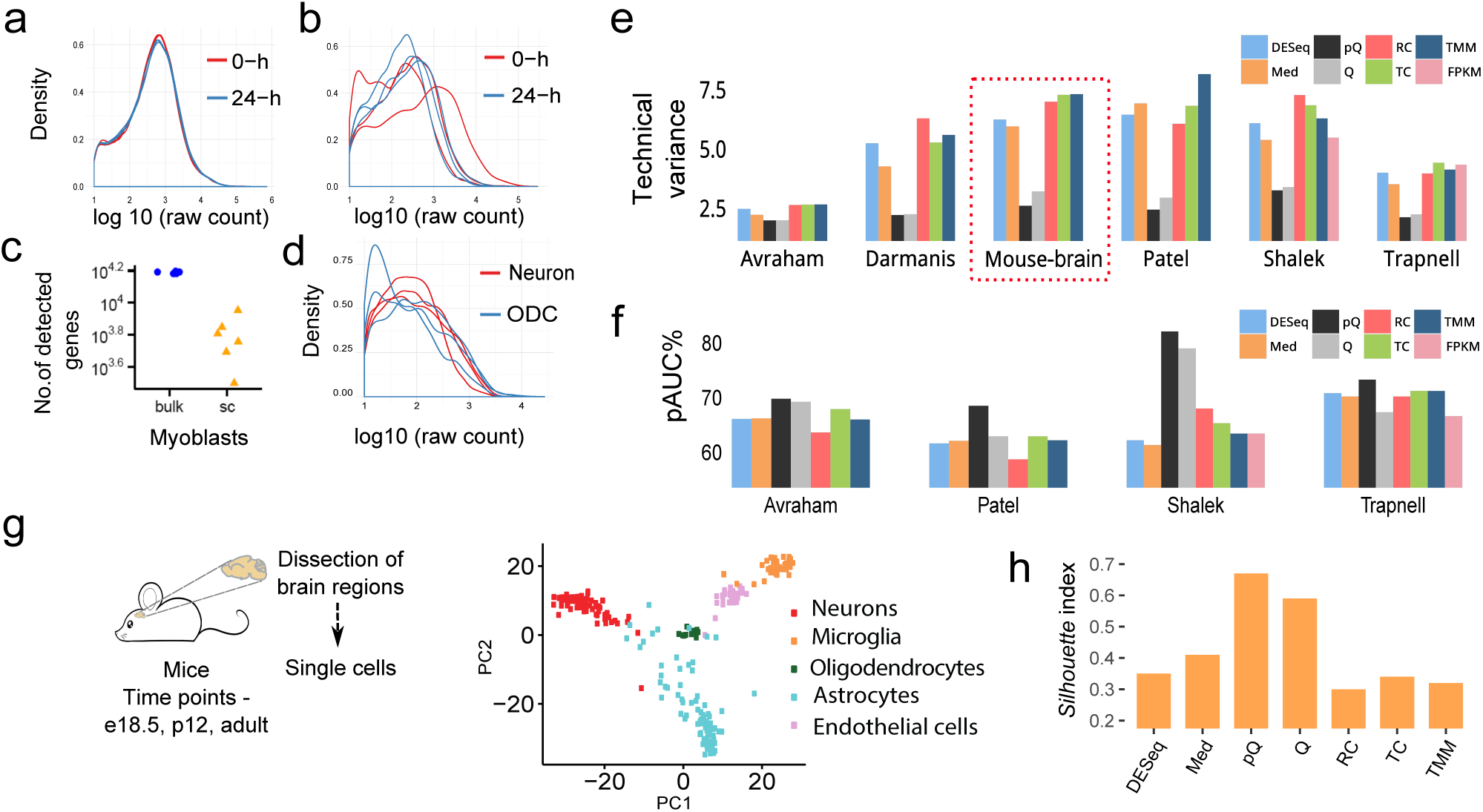
Comparison of single-cell RNA-seq normalization methods. (a) Gene expression distribution (raw read counts ≥10) in myoblast bulk RNA-seq: 3 replicates from each condition. Since the six distributions are nearly identical, only two are visible when overlaid. (b) Distribution of gene expression values in single myoblast cells before and after differentiation: 3 randomly chosen cells from each condition. (c) Number of detected genes in the 6 bulk samples and 6 singlecells from (a) and (b), respectively. (d) Distribution of gene expression values in randomly chosen postmitotic single cells from adult human cerebral cortex: 3 oligodendrocytes (ODC) and 3 neurons.^23^ (e) Percent of expression variance in single-cell RNA-seq data that is attributable to technical bias *b* for the raw counts (8 normalization methods,6 datasets). (f) Accuracy of single-cell DE calls on four scRNA-seq datasets, quantified using partial-AUC (pAUC) of receiver operating characteristic (ROC).^24^ Accuracy is defined as concordance with bulk-sample DE genes. (g) Principal component analysis (PCA) of mouse brain single cell transcriptomes (GSE79374) of the five most prevalent cell types (neurons, microglia, oligodendrocytes, astrocytes, myelinating oligodendrocytes and endothelial cells), after pQ normalization. (h) *Silhouette* index (cell type separability score) of mouse brain transcriptomes after expression normalization by multiple methods.

Strategies originally devised for bulk RNA-seq are also commonly employed for identifying DE genes in scRNA-seq data.In particular, previous single-cell studies assumed that gene expression variation followed a well-defined parametric distribution.^6,8,9^ Parametric tests were therefore used to estimate the statistical significance of differentialexpression. However, it is well known that when the number of samples is large, as is typical of single-cell data, nonparametric tests have very similar statistical power.^6^ In fact, when the true distribution violates parametric assumptions, parametric methods are actually less powerful.^10^ Parametric DE algorithms are also typically slower, due to greater computational complexity. Parametric tests are therefore not ideally suited to data from many hundreds or thousands of single cells.^4^ Thus, it is important to develop nonparametric methods for single-cell DE analysis.

We propose a two-pronged approach to address the above problems, which can be applied to any single cell omics dataset because it does not require the expression distribution to have a specific form. 1) For expression normalization, we use a variant of quantile (Q) normalization.^11^ One drawback of Q normalization is that it does not address variability in the number of detected genes. Consequently, if a cell type A has fewer detected genes than another cell type B, many lowly expressed genes will remain undetected in the former, resulting in artifactual DE calls. Scaling-based normalization methods are also susceptible to this artifact. Our approach, which we term pseudocounted Quantile (pQ) normalization, homogenizes the expression of all genes below a fixed rank in each cell (Supplementary Material). 2) For DE analysis, we introduce a novel nonparametric statistic that combines rank-based and mean-based measures of expression divergence.

Methods for scRNA-seq normalization have not previously been systematically evaluated. For comparative benchmarking, we accessed scRNA-seq data from four previous studies that also performed bulk RNA-seq on the same samples^12–15^ (Supplementary Table 1). In addition to pQ normalization, we tested seven existing normalization methods: Q, total read count^16^ (TC), median^16^ (Med), DESeq, TMM, FPKM and raw read count (RC) quantified using HTSeq^17^ (Count; Supplementary Fig. 1). Expression was quantified as normalized read counts in all cases, except in the case of FPKM normalization. Performance was evaluated based on three criteria: 1) proportion of expression variance attributable to technical bias, 2) accuracy of DE calls and 3) transcriptomic separability of cell types.

It has recently been reported that a substantial proportion of scRNA-seq expression variability is explained by a single technical factor: the number of detected genes (b) in each cell.^18^ A good normalization method should minimize the impact of this technical bias. We, therefore, calculated the post-normalization expression variance explained by b, and used it as a measure of normalization efficacy (Supplementary Material). We performed this analysis on five published scRNA-seq datasets namely Avraham, Darmanis, Patel, Shalek and Trapnell (Supplementary Table 1). In addition, we generated a sixth scRNA-seq dataset consisting of 333 single cells from mouse brain, isolated at three developmental time points from three brain regions (Supplementary Material). The latter dataset provides a rigorous in vivo benchmark representing multiple co-existing cell types processed in multiple batches. On all six datasets, including mouse brain, pQ normalization was the most effective in minimizing technical bias, closely followed by conventional Q normalization (Fig. 1e).

An accurate normalization method should also improve the quality of subsequent DE gene analysis. For each of the 7 count-based normalization methods, we therefore examined the accuracy of single-cell DE gene calls (we tested the performance of FPKM normalization only for Shalek and Trapnell datasets). A single-cell DE gene call was counted as a true positive if the gene was also present in the matched bulk-transcriptome DE list. If not, it was counted as a false positive. The Wilcoxon rank sum test^19^ was used to call single-cell and bulk DE genes. This analysis was performed on the four datasets that contained matched single-cell and bulk profiles (Avraham, Patel, Shalek, Trapnell). On all four datasets, pQ normalization maximized the accuracy of single-cell DE analysis (Fig. 1f; Supplementary Fig. 2). TC and Q were second and third respectively, based on average rank (Supplementary Table 2).

In order to devise a nonparametric test for single-cell DE analysis, we first considered the Wilcoxon rank sum test.^19^ One potential limitation we anticipated was that this rank-based test ignored the magnitude of expression deviation between the two groups of cells. We therefore also calculated the expression-difference statistic D from the NOISeqBIO algorithm^20,21^ (Supplementary Material). The D-statistic exploits the fact that, under the central limit theorem, the distribution of the mean expression value in each cell population is approximately Gaussian. It is thus broadly applicable, regardless of the parametric form of the underlying single-cell expression distributions. It is also robust since outliers are removed by the median-based pQ-normalization step. We then used Fisher’s method to combine p-values from the two statistical tests into a single, composite p-value for differential expression.

The DE analysis method described above, which we named <u>NO</u> nparametric <u>D</u> ifferential <u>E</u> xpression for <u>S</u> ingle-cells (NODES), was compared to three methods originally designed for bulk-sample DE analysis (DESeq2,^22^ edgeR^8^ and NOISeqBIO^10^), one recently developed algorithm for single-cell DE analysis (scde^9^) and also the Wilcoxon rank sum test (the D-statistic, by itself, was not found to be effective). As above, we applied these methods to the Avraham, Patel, Shalek and Trapnell datasets and used the corresponding bulk-sample DE gene calls for benchmarking. To facilitate uniform comparison, all DE algorithms were provided with the same TMM-normalized expression matrix as input. On all four datasets, NODES generated the most accurate DE calls (Fig. 2a, Supplementary Fig. 6) and the Wilcoxon test was on average second-best (Supplementary Table 3). The same was true when the DE algorithms were allowed to use their own default normalization strategies. We also tested the DE analysis algorithms on 10 simulated datasets generated by resampling expression values from a genuine scRNA-seq dataset (Supplementary Material). Again, NODES clearly outperformed the other methods(Supplementary Fig. 7). As a resource, we therefore used NODES to define marker genes uniquely expressed in each of the five major mouse brain cell types (Fig. 2b; Supplementary File).

**Figure 2.**
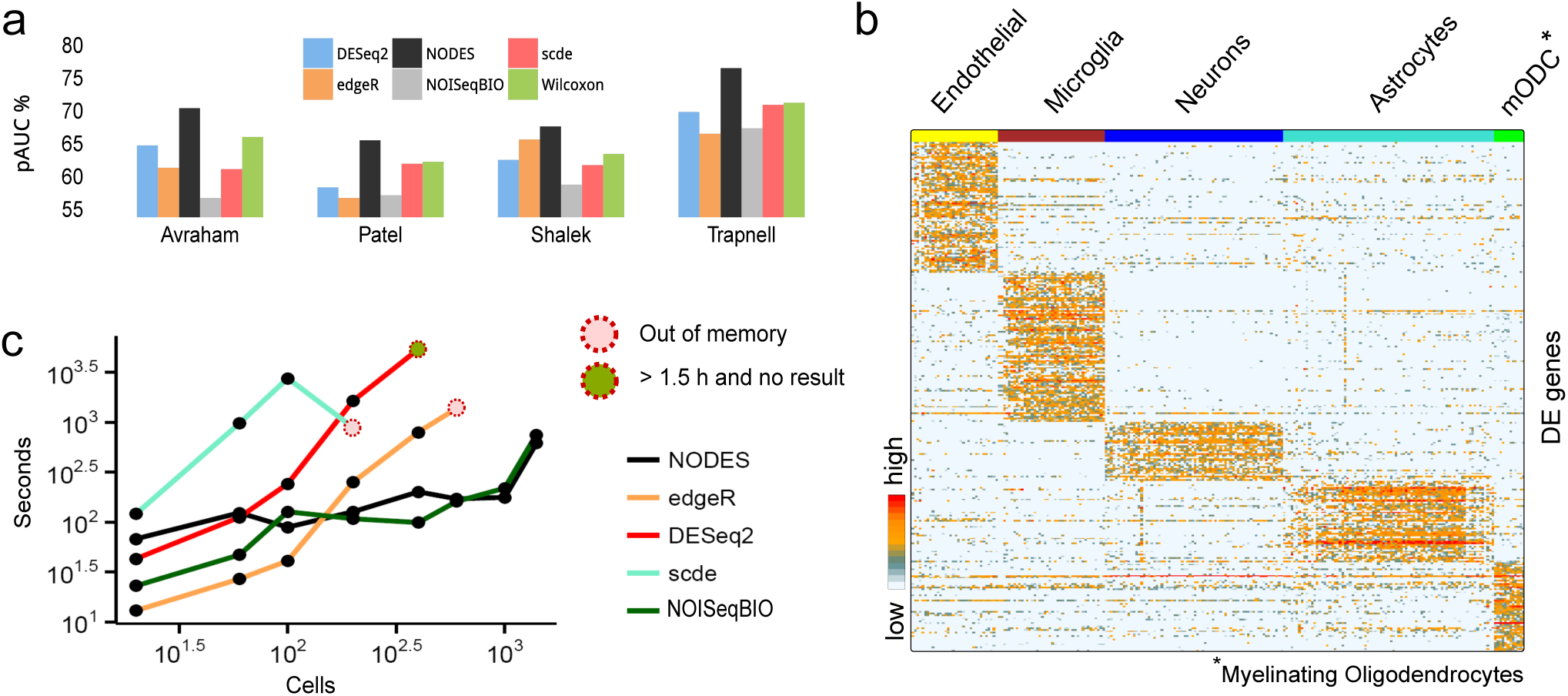
Comparison of methods for calling single-cell DE genes. (a) Partial-AUC (pAUC) estimates of single-cell DE calls, with bulk DE genes as benchmark. (b) Expression pattern shown for cell-type specific genes for the 5 major cell types depicted in Fig. 1g, determined by NODES on pQ normalized expression data. (c) Execution time of DE analysis programs on scRNA-seq data from mouse ES cells before and after LIF withdrawal (inDrop/ Drop-seq; Klein data). X-axis: total number of analyzed cells. Equal numbers of cells were selected from each condition.

Given the high level of noise in scRNA-seq data, large numbers of cells are increasingly being profiled in a single experiment in order to achieve sufficient statistical power for downstream analyses. Scalability is thus an essential feature of scRNA-seq analysis algorithms. We used a collection of 1,596 scRNA-seq datasets^4^ (inDrop/ Drop-seq protocol) to measure execution time for the five tested DE analysis algorithms (Supplementary Material). Notably, the parametric methods (scde, DESeq2 and edgeR) could not easily scale beyond 200-400 cells, either due to memory limitations or excessive run time (Fig. 2c). In contrast, the non-parametric methods produced results relatively quickly, even on datasets as large as ~ 1,600 cells. The above results demonstrate the value of pQ normalization in reducing technical variability and improving the quality of downstream expression analysis. They also highlight the superior accuracy of NODES in calling DE genes, relative to existing methods. One significant advantage of the nonparametric methods introduced in this study is that they can straightforwardly be applied to scRNA-seq data from any experimental protocol, since they make no assumptions about the shape of the distribution. The nonparametric DE approach also provides a transformative reduction in computational complexity and execution time, which will be crucial for analyzing the massive single-cell datasets generated by inDrop/ Drop-seq and other high-throughput single cell technologies.

## Software

R package implementing pQ and NODES can be found at https://goo.gl/Ndx07M.

## Acknowledgments

This work is supported by grant #SPF2012/003 from the Agency for Science, Technology and Research (A*STAR), Singapore.

## Author Contributions

DS, SP and BL conceived the study. DS and SP designed the statistical methods and developed the analysis strategies. DS implemented the methods and performed the analyses. NAR performed the single cell RNA-seq experiments with assistance from ML. DS and SP wrote the manuscript with help from BL and NAR. All authors read and approved the manuscript.

## References

1. Shapiro, E., Biezuner, T. & Linnarsson, S. Single-cell sequencing-based technologies will revolutionize whole-organism science. Nature Reviews Genetics 14, 618–630 (2013).

2. Saliba, A.-E., Westermann, A. J., Gorski, S. A. & Vogel, J. Single-cell rna-seq: advances and future challenges. Nucleic acids research gku555 (2014).

3. Kim, J. K., Kolodziejczyk, A. A., Illicic, T., Teichmann, S. A. & Marioni, J. C. Characterizing noise structure in single-cell rna-seq distinguishes genuine from technical stochastic allelic expression. Nature communications 6 (2015).

4. Klein, A. M. et al. Droplet barcoding for single-cell transcriptomics applied to embryonic stem cells. Cell 161, 1187–1201 (2015).

5. Trapnell, C. et al. Transcript assembly and quantification by rna-seq reveals unannotated transcripts and isoform switching during cell differentiation. Nature biotechnology 28, 511–515 (2010).

6. Anders, S. & Huber, W. Differential expression analysis for sequence count data. Genome biol 11, R106 (2010).

7. Robinson, M. D., Oshlack, A. et al. A scaling normalization method for differential expression analysis of rna-seq data. Genome Biol 11, R25 (2010).

8. Robinson, M. D., McCarthy, D. J. & Smyth, G. K. edger: a bioconductor package for differential expression analysis of digital gene expression data. Bioinformatics 26, 139–140 (2010).

9. Kharchenko, P. V., Silberstein, L. & Scadden, D. T. Bayesian approach to single-cell differential expression analysis. Nature methods11, 740–742 (2014).

10. Tarazona, S. et al. Data quality aware analysis of differential expression in rna-seq with noiseq r/bioc package. Nucleic acids research gkv711 (2015).

11. Bolstad, B. M., Irizarry, R. A., AÅstrand, M. & Speed, T. P. A comparison of normalization methods for high density oligonucleotide array data based on variance and bias. Bioinformatics 19, 185–193 (2003).

12. Avraham, R. et al. Pathogen cell-to-cell variability drives heterogeneity in host immune responses. Cell 162, 1309–1321 (2015).

13. Patel, A. P. et al. Single-cell rna-seq highlights intratumoral heterogeneity in primary glioblastoma. Science 344, 1396–1401 (2014).

14. Shalek, A. K. et al. Single-cell rna-seq reveals dynamic paracrine control of cellular variation. Nature (2014).

15. Trapnell, C. et al. Pseudo-temporal ordering of individual cells reveals dynamics and regulators of cell fate decisions. Nature biotechnology 32, 381 (2014).

16. Dillies, M.-A. et al. A comprehensive evaluation of normalization methods for illumina high-throughput rna sequencing data analysis. Briefings in bioinformatics 14, 671–683 (2013).

17. Anders, S., Pyl, P. T. & Huber, W. Htseq–a python framework to work with high-throughput sequencing data. Bioinformatics btu638 (2014).

18. Hicks, S. C., Teng, M. & Irizarry, R. A. On the widespread and critical impact of systematic bias and batch effects in single-cell rna-seq data. bioRxiv 025528 (2015).

19. Wilcoxon, F. Individual comparisons by ranking methods. Biometrics bulletin 80–83 (1945).

20. Tarazona, S., GarcÍa-Alcalde, F., Dopazo, J., Ferrer, A. & Conesa, A. Differential expression in rna-seq: a matter of depth. Genome research 21, 2213–2223 (2011).

21. Efron, B., Tibshirani, R., Storey, J. D. & Tusher, V. Empirical bayes analysis of a microarray experiment. Journal of the American statistical association 96, 1151–1160 (2001).

22. Love, M. I., Huber, W. & Anders, S. Moderated estimation of fold change and dispersion for rna-seq data with deseq2. Genome Biol 15, 550 (2014).

23. Darmanis, S. et al. A survey of human brain transcriptome diversity at the single cell level. Proceedings of the National Academy of Sciences 112, 7285–7290 (2015).

24. Robin, X. et al. proc: an open-source package for r and s+ to analyze and compare roc curves. BMC bioinformatics 12, 77 (2011).

